# Gut microbial metabolic flux disorder in hypertension

**DOI:** 10.1101/2024.10.09.617349

**Authors:** Wenkai Lai, Yuchen Zhang, Meiling Wang, Shirong Lai, Qing Liu, Qi Luo, Quan Zou, Fenglong Yang

**Affiliations:** Department of Bioinformatics, Fujian Key Laboratory of Medical Bioinformatics, School of Medical Technology and Engineering, Fujian Medical University, Fuzhou, China; Institute of Fundamental and Frontier Sciences, University of Electronic Science and Technology of China, Chengdu, China

**Keywords:** Hypertension, microbes, gut, metabolites

## Abstract

Hypertension is a major risk factor for cardiovascular diseases such as stroke and heart failure. Recent studies have shown that changes in the composition and function of the gut microbiota are closely related to the onset and development of hypertension. However, the individual differences in gut microbiota species make it difficult for traditional analysis methods to effectively reveal the pathogenic mechanisms of hypertension. In contrast, the inter-individual variability in gut microbial metabolites is much smaller, allowing for better cross-individual comparisons and reducing confounding factors in analysis. The interactions between gut microbiota and metabolites are highly complex, and network analysis can systematically capture this complexity. In this study Flux Balance Analysis (FBA) was utilized to predict the metabolic flux of gut microbiota and constructed cross-feeding networks. Random Forest and XGBoost models were employed to identify metabolites associated with hypertension. A differential microbial correlation network was used to analyze important metabolically related microbial sub-networks, and ultimately, the metabolic abnormalities and metabolite-related pathways were analyzed at the network level using the metabolite correlation network and cross-feeding networks. It was observed that the interaction patterns among 25 species—collectively referred to as the KEPR guild, with the most abundant genera being Eubacterium, Ruminococcus, Klebsiella, and Parabacteroides—changed, leading to alterations in 12 metabolites, such as choline (chol), 1-butanol (btoh), trimethylamine (tma), cytidine (cytd), and betaine (glyb) etc. Choline can be oxidized to form betaine, thereby affecting host blood pressure. Abnormalities in siroheme and methanethiol may result in reduced secretion of hydrogen sulfide by microbes, which in turn impacts blood pressure regulation mechanisms. The changes in these 12 metabolites may also enhance the degradation of mucin-type O-glycans and reduce butyrate metabolic activity, weakening the protective ability of intestinal epithelial cells. This may lead to inflammation and oxidative stress, exacerbating endothelial cell damage and consequently resulting in endothelial dysfunction and increased blood pressure. The findings of this study provide new insights into the pathogenic mechanisms of hypertension and offer potential targets for clinical intervention.

## 1 INTRODUCTION

Hypertension remains one of the major risk factors for cardiovascular diseases (CVD) such as stroke and heart failure. In addition, it is an important associated factor for common comorbidities such as chronic kidney disease, obesity, and type 2 diabetes^1,2^. In 2010, approximately 31% of the global population had hypertension, making it a global public health issue^3^. Despite extensive research and interventions, blood pressure control still faces many challenges^4^.

Recent studies have shown that the gut microbiota plays a crucial role in the development of cardiovascular diseases^5^. For instance, patients with heart failure with preserved ejection fraction (HFpEF) exhibit significant gut dysbiosis, with marked differences compared to healthy individuals^6^. The gut microbiota metabolizes dietary choline, phosphatidylcholine, and L-carnitine to produce trimethylamine (TMA), which is further oxidized to trimethylamine N-oxide (TMAO), a metabolite known to promote atherosclerosis^5,7^. These findings provide new perspectives and insights into the etiology of hypertension and potential therapeutic strategies.

The gastrointestinal tract is the largest immune cell compartment in the body, representing the intersection between the environment and the host^8^. Lifestyle can shape and be influenced by the microbiome^9–11^, thereby altering the risk of developing hypertension. A well-studied example is the consumption of dietary fiber, which leads to the production of short-chain fatty acids and contributes to the expansion of anti-inflammatory immune cells, thus preventing the progression of hypertension^12^.

The gastrointestinal tract is also the primary site for the interactions between dietary components, gut microbiota and their metabolites, and antihypertensive drugs, with blood pressure potentially regulated by gastrointestinal hormones^13^.

Although species-level analysis can provide basic information about microbial composition, it does not fully capture the complexity and functional diversity of microbial communities. Species-level analysis overlooks interactions between microorganisms and their functional potential. Currently, research on hypertension-related gut microbiota is relatively limited. Traditional microbial studies often focus on changes in microbial species abundance, such as *α*-diversity, *β*-diversity, and species abundance differences^14^. However, challenges in microbial research include data sparsity and significant inter-individual variability, which complicate and reduce the accuracy of constructing risk assessment or sample classification models. The interaction networks between microorganisms are crucial for maintaining gut health and metabolic balance^15,16^, yet these interactions may not be fully captured at the species level. The impact of gut microbiota on the host is typically mediated through interactions between their metabolites and the host.^17,18^. Compared to simple species analysis, metabolomics has many advantages in deciphering the composition and function of microbes within an organism: Gut microbiota metabolites, as signaling molecules and substrates for host metabolic responses, influence various physiological and pathological processes in the host^19^. For example, short-chain fatty acids (SCFAs) are one of the most studied classes of small molecule metabolites. They are produced by gut microbes through the fermentation of dietary fibers and can regulate host physiological and biochemical functions. These functions include maintaining the natural intestinal barrier at the colonic epithelium and mucus levels, regulating gut motility, secreting gut hormones, modulating chromatin, influencing the gut-brain axis, and supporting immune functions. Metabolites are the end products of metabolic activities in the body, and their composition and abundance directly reflect the physiological state, metabolic pathways, and levels of biological activity within the organism^20,21^. Metabolomics also offers strong biological interpretability. Metabolites are often closely associated with physiological processes and metabolic pathways in the body^22^, which endows metabolomic data with significant biological relevance and interpretability.

Recent studies have shown that hypertension is closely related to disruptions in gut microbiota metabolites^23^. The gut microbiota can influence host health, particularly through its metabolites. These metabolites can enter the bloodstream and affect the host’s physiological and pathological states. For example, some studies have found that certain gut microbial metabolites, such as short-chain fatty acids (SCFAs), can counteract hypertension by modulating immune responses, reducing inflammation levels, regulating energy balance, and cholesterol metabolism^24^. Additionally, some gut microbiota-derived metabolites may be involved in blood pressure regulation, such as trimethylamine N-oxide (TMAO) produced from dietary nutrients by gut microbes, which has been associated with poor cardiovascular health, including atherosclerosis, hypertension, and heart disease^7^.

In this study, we constructed individualized whole-body metabolic models and calculated the metabolic flux spectrum for each sample through metabolic flux analysis. Metabolic flux reflects the rate at which microorganisms produce and consume metabolites, which is related to the disease state of the host. This research focuses on metabolites to explore the relationship between metabolites and disease phenotypes, ultimately pinpointing specific species. We systematically organized microbiome data for comprehensive analysis, constructing a gut microbial interaction network(GMiN), a metabolic flux interaction network(GMFiN) based on statistical methods, and a knowledge-based microbial cross-feeding network(GMxN). These networks were used to calculate network metrics such as degree centrality, betweenness centrality, closeness centrality, and eigenvector centrality of the network nodes, as well as to assess differences in edges between nodes. This enabled a systematic analysis of the interactions between microbes and metabolites and their relationship with the occurrence of hypertension.

## 2 METHODS

### 2.1 Data Collection and Processing

In this study, hypertension data from three studies were collected as the discovery cohort from the curatedMetagenomicData databases^25^. To precisely investigate gut metabolic dysregulation in hypertension, we only analyzed samples from individuals with hypertension who had no other diseases. Among these samples, 169 met the criteria, but some lacked age records, with the overall age distribution ranging from 55 to 70 years. Therefore, the criteria for selecting healthy samples were an age between 55 and 70 years and a BMI between 18.5 and 23.9, resulting in the collection of 109 healthy samples.

Additionally, 441 samples were collected from the study by Jacobo de la Cuesta-Zuluaga etal.^26^, with each sample providing detailed clinical information, including age, gender, BMI, diastolic pressure, systolic pressure, cholesterol, LDL, cardiometabolic diseases, and medication use. To extract samples in fully healthy and fully hypertensive states, we considered not only diastolic and systolic pressures but also age, BMI, cholesterol levels, cardiometabolic diseases, and medication use. Since the ages of the samples in this dataset are relatively younger, we selected healthy samples with an age greater than 35 years, a BMI between 18.5 and 25, a diastolic pressure below 80 mmHg, a systolic pressure below 130 mmHg, and cholesterol below 200 mg/dL. For hypertensive samples, we selected those older than 35 years with a diastolic pressure above 90 mmHg, a systolic pressure above 140 mmHg, and no history of medication use. Ultimately, 41 samples were collected, including 19 healthy and 22 hypertensive samples (Figure 1).

**Figure 1.**
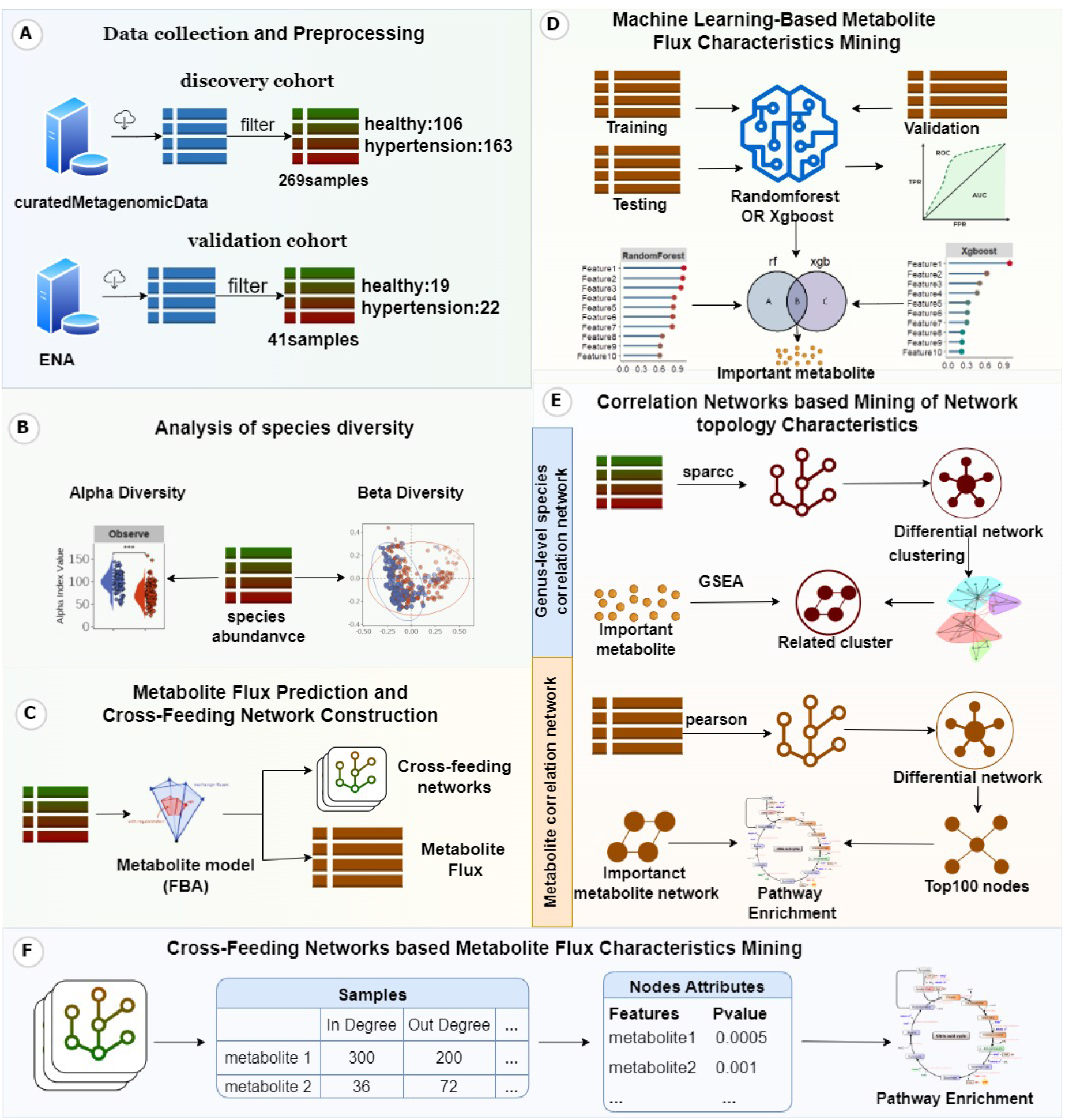
Research workflow for studying the mechanisms of dysregulated gut microbiota metabolic flux in hypertension: Downloading and preprocessing of hypertension gut microbiota data (Step A), analysis of microbial species diversity (Step B), prediction of metabolic flux based on microbial data (Step C), construction of a hypertension diagnostic model using metabolic flux data, and calculation of important features in the model (Step D). Then calculate the differential microbial correlation network and enrich the subnetworks of the differential microbial correlation network using the important features. Calculating the differential metabolite correlation network and the important feature-related subnetworks, followed by pathway enrichment analysis of differential nodes and important feature subnetwork nodes (Step E). Constructing individualized cross-feeding networks, calculate the topological features of nodes in each network, conduct differential analysis of each node, and perform pathway enrichment analysis (Step F).

Based on the microbial metabolic reconstruction model database AGORA1 developed by Ines Thiele etal.^27^, we mapped the individualized gut microbiota species abundance profiles to this database. Since the database contains 818 species, we retained only the abundance profiles of these 818 species. In some samples, the abundance of these species was zero, making them irrelevant for model construction; therefore, we only retained samples with a relative abundance greater than 60%. Finally, 269 samples remained in the discovery cohort, including 106 healthy and 163 hypertensive samples. The validation cohort consisted of 41 samples, including 19 healthy and 22 hypertensive samples.

### 2.2 Analysis of Gut Microbiota Diversity Differences Between Hypertensive and Healthy Populations

#### 2.2.1 *α****-Diversity Analysis***

To explore the relationship between species diversity and hypertension, six *α*-diversity indices of gut microbiota were calculated using the R package MicrobiotaProcess^28^. These indices reflect the diversity and evenness of species within a habitat or community by calculating the composition and abundance of species within samples^29,30^. And then the Wilcoxon test was used to compare the significance of microbial alpha diversity between the healthy and hypertensive groups.

### 2.2.2 *β****-Diversity Analysis***

To analyze the overall differences in gut microbiota structure between hypertensive and healthy populations. The PCoA method was used to illustrate the relationships between different groups. First, we calculated the distances between samples using the Bray-Curtis method^31^and evaluated the degree of difference between the two groups using the PERMANOVA method^32^.

### 2.3 Analysis of Differences in Gut Microbiota Metabolite Flux Between Hypertensive and Healthy Populations

#### 2.3.1 Gut Microbiota Metabolic Flux Prediction

In this experiment, we constructed a database containing metabolic information from multiple species and then mapped the species abundance profiles of the samples to this database. Using the COBRA method^33^, we constructed a metabolic model for each individual sample. Dietary data were further incorporated as constraints, and metabolic flux spectra for each sample were obtained using Flux Balance Analysis (FBA)^34^. Using metabolic flux data can avoid the issue of data sparsity commonly associated with species-level analysis, while also addressing challenges related to metabolite profiling. By extracting metabolic flux features, it is possible to trace back to the species level, thereby providing more precise clinical recommendations.

### 2.3.2 Construction of a Hypertension Diagnosis Model and Metabolic Importance Feature Extraction

To explore the differences in metabolites between the two sample groups, traditional differential analysis methods or linear models struggle to accurately identify important features due to the complexity of microbial and microbial reaction data. Therefore, this study employed two rule-based tree structure models, Random Forest and XGBoost, to perform binary classification modeling of the metabolic flux data, classifying samples into healthy and hypertensive groups, and identifying important features.

First, the samples in the discovery cohort were divided into a training set and a validation set in a 7:3 ratio. The Random Forest and XGBoost models were trained using the training set, and model performance was optimized using data from the validation set. Model parameters were updated to avoid overfitting. The test cohort was used as an independent validation set to further assess model performance. However, due to differences in age distribution between the test cohort and the discovery cohort, as well as differences in the countries where the samples were sequenced, the test cohort data were only used for validating the machine learning models. Once the model performance met the expected criteria, we extracted the important features from the models. To ensure comparability of the importance scores between the two models, the importance scores calculated by both models were normalized (formula 1). We then selected the intersection of the top 20 important features from both models. The importance scores of the intersected features were determined by multiplying the normalized scores of both models by the AUC values of the models on the test set as weights (formula 2).

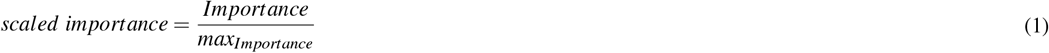

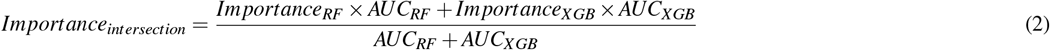

### 2.4 Networks based Mining of Network topology Characteristics

#### 2.4.1 Microbial Correlation Differential Network Analysis

The gut microbiome is a complex system. Recent studies have shown that hypertension is closely related to disruptions in the metabolism of gut microbiota. Gut microbiota can influence host health, particularly through their metabolic products. Since gut microorganisms often function cooperatively, we used the microbial species abundance matrix to calculate the correlations between species using the SparCC method. We then constructed a microbial correlation network by sparsifying the matrix using the overall mean correlation as a threshold. To calculate the microbial correlation differential network, we subtracted the edge weights of the healthy microbial network from those of the hypertensive microbial network. We then used the permutation test^35^ to assess the significance of the edge differences between the healthy and hypertensive networks, filtering out non-significant edges. This allowed us to compute the differential microbial correlation network.

### 2.4.2 Clustering and Enrichment of Microbial Nodes in the Differential Network Related to Metabolic Important Features

To explore which microorganisms mainly contribute to significant metabolic fluxes, this experiment calculated the differential correlation network of microorganisms and used a random walk algorithm^36^ to cluster the nodes. The enrichment score of each guild for the metabolic important features was calculated using the GSEA algorithm^37,38^.

First, this study used the cross-feeding network of each sample to statistically count the number of samples in which an absorption or secretion relationship exists between metabolic important features and each species. For example, if species A was found to secrete metabolite A in 100 samples, then the sample count for species A secreting metabolic important feature 1 is 100. If species A was found to secrete metabolic important feature 2 in 80 samples, then the sample count for species A secreting metabolic important feature 2 is 80. In this way, the total count for species A would be 180. The species were then ranked based on their counts. Finally, the distribution of the ranks within each guild was used to calculate the enrichment score for each guild concerning the important features, along with the statistical significance.

### 2.5 Differential Network Analysis of Hypertension-Related Metabolic Flux Correlations

Metabolites can form metabolic pathways through interactions, and analyzing the network topology and edges can help us understand the relationship between hypertension and the gut more precisely. In this study, we used the metabolic flux matrix to calculate the metabolite correlation networks for healthy and hypertensive samples using the pearson correlation method. To compute the differential metabolite correlation network, we performed a significance difference analysis on the edges using Fisher’s z-test^39^. The edge weights of the hypertensive metabolite network were subtracted from those of the healthy network, and non-significant edges were filtered out to obtain the differential metabolite correlation network.

We calculated the topological features of each node in both networks, such as Betweenness, Closeness, Degree, and Eigen centrality. To identify nodes with the most significant differences in topological features, we extracted the top 20 nodes with the largest differences in topological features between the two networks. We then used the KS-test to calculate the significance of the top 20 nodes with different topological features between the healthy and hypertensive networks.

### 2.6 Cross-Feeding Network Analysis of Gut Microbiota

In this study, the metabolic flux spectrum for each sample was calculated using FBA, allowing us to determine the rate at which each species produces and consumes metabolites within each sample. This enabled the construction of a network for each sample, with species and metabolites serving as nodes. Since this study focuses more on the interactions between metabolites and samples, we calculated the topological feature values of the metabolites within each network. The Wilcoxon test was used to analyze the significance of differences in the topological features of metabolite nodes between the two groups, thereby identifying metabolites with significant differences.

### 2.7 Pathway Enrichment of Differential Network Nodes

To explore the pathways and functions involved with these metabolites, pathway enrichment analysis was performed on the metabolites of interest. For GMFiN network node enrichment, this study first separated nodes with upregulated and downregulated degrees. We then gathered microbial-related reaction equations from the VMH database, using the reactants and products of these equations to identify metabolites associated with specific pathways. Enrichment of nodes within these pathways was performed using the cumulative hypergeometric distribution method. In the GMxN, where the metabolite production rate by microbes allows for a more precise understanding of microbial metabolic pathways, we similarly separated nodes with upregulated and downregulated indegrees. Here, products from the reaction equations were treated as pathway-related metabolites, and pathway enrichment of the nodes was also conducted using the cumulative hypergeometric distribution method.

## 3 RESULTS

### 3.1 Gut Microbial Diversity were Reduced in Hypertensive Patients

In this study, gut microbial data were divided into two groups based on disease status: hypertensive and healthy. We conducted *α*-diversity analysis on the gut microbiota of both the hypertensive and healthy groups, calculating six indices: Observe, Chao1, ACE, Shannon, Simpson, and Pielou. In the discovery cohort, five of these indices showed significant differences, with the healthy group exhibiting higher values than the hypertensive group (Figure 2A), indicating a decrease in microbial richness and evenness in hypertensive patients.

**Figure 2.**
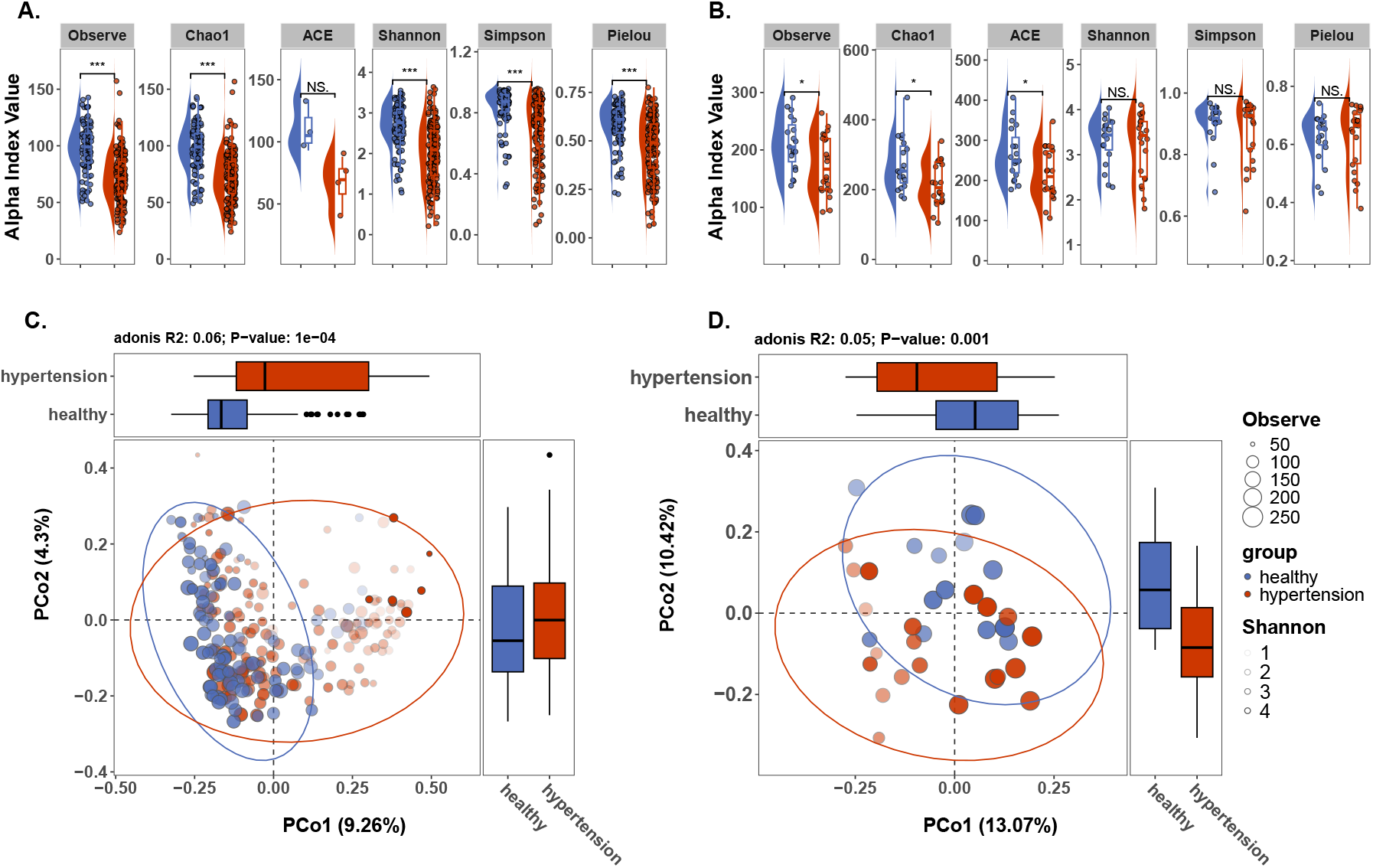
Gut Microbial Diversity were Reduced in Hypertensive Patients: five of these indices Observe, Chao1, Shannon, Simpson and Plelou reduced in Hypertensive Patients from discovery cohort (A). while only Observe, Chao1 and ACE reduced in Hypertensive Patients from validation cohort(B). significant gut microbial composition differences between the hypertensive and healthy samples were detected from discovery cohort (C) and validation cohort (D).

The *β*-diversity analysis revealed that the inter-group distances between healthy and hypertensive samples were greater than the intra-group distances. A permutational multivariate analysis of variance (PERMANOVA) test showed an R-squared value of 0.05 and a p-value of 0.001 (Figure 2C), indicating significant differences between the hypertensive and healthy samples. In the validation cohort, the differences in *α*-diversity were not as pronounced as in the discovery cohort, with only three indices showing significant differences, and again, the hypertensive group exhibited lower diversity compared to the healthy group (Figure 2B). The *β*-diversity analysis results were consistent with those of the discovery cohort (Figure 2D).

Although the results from the two datasets were generally similar, the three diversity indices that did not match could be attributed to differences in the age distribution between the two cohorts. Additionally, gut microbiota varies significantly between individuals, and microbial data is often sparse, leading to differences in microbial composition across samples.

### 3.2 Gut microbial metabolic patterns are altered in patients with hypertension

Analyzing species abundance data alone cannot fully capture the complexity and functional variability of microbial communities, and it may overlook interactions and functional potential among microbes. Microbes interact with the host primarily through metabolites, which exhibit less variability between samples compared to species abundance data(Figure 3A). Therefore, investigating the pathogenic mechanisms of hypertension by focusing on metabolites is likely to yield more insightful results. In this study, we utilized Random Forest and XGBoost models to identify key features, finding that the AUC reached 0.87 for the Random Forest model and 0.93 for the XGBoost model in the testing set. In the validation set, the AUC for the Random Forest model was 0.84, and 0.8 for the XGBoost model(Figure 3B). We then identified the top 20 important features for both models(Figure 3C) and found an intersection of 12 key features. Of these, 8 were related to species-environment exchange reactions (extracellular space), and 4 involved community-gut lumen exchange reactions (medium)(Figure 3D).

**Figure 3.**
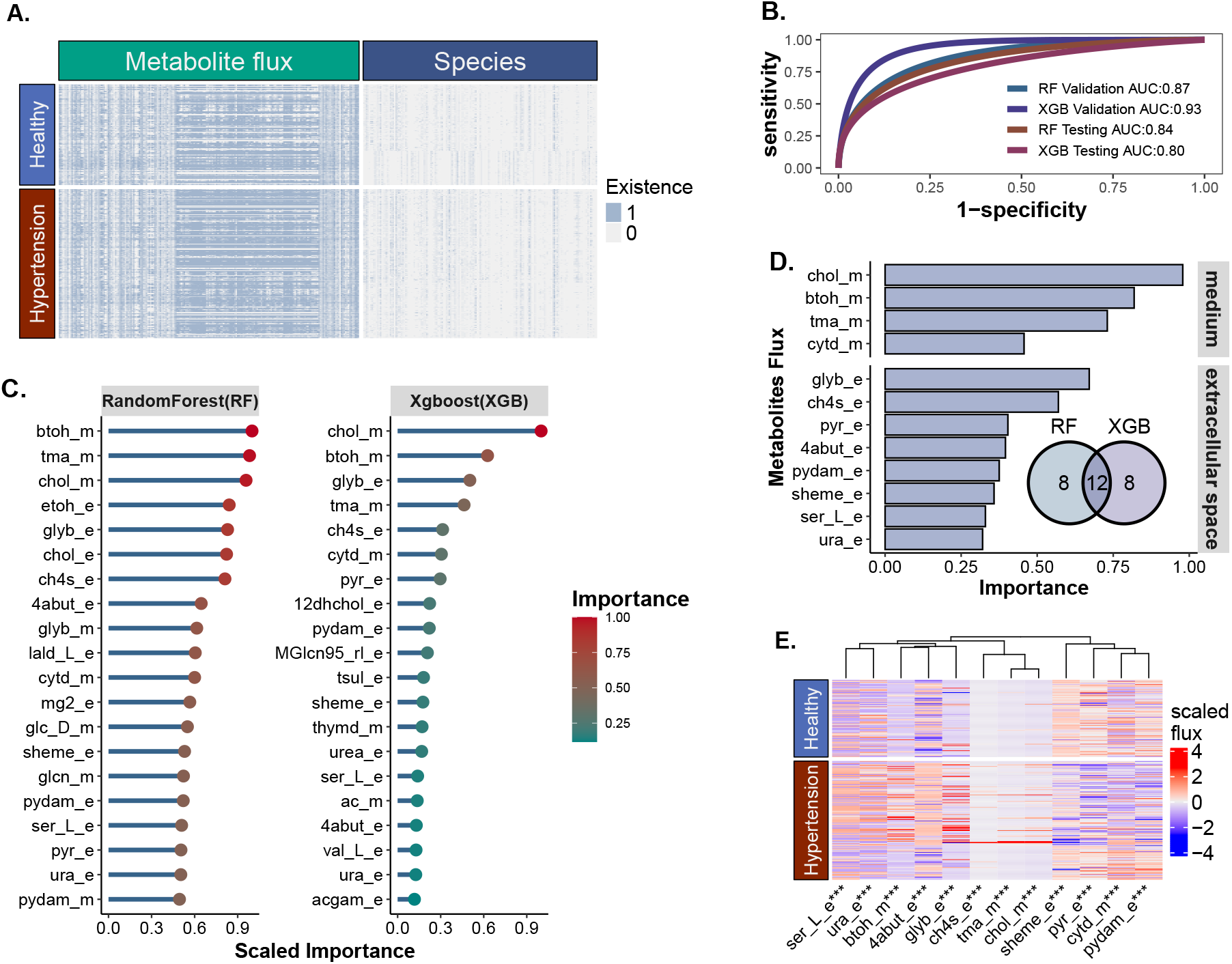
Gut microbial metabolic patterns are altered in patients with hypertension: Metabolite flux data is not as sparse and sample-specific as species data. In the heatmap, blue represents the detection of a species or metabolite, while gray indicates its absence (A). The performance of the Random Forest and XGBoost models constructed using metabolite flux data for classifying healthy and hypertensive samples is excellent. The models were evaluated on the validation and test sets using ROC curves (B). The important features identified by both models, with their importance scores normalized (C). Both models identified 12 important metabolite. The importance scores were weighted by the AUC values of the models on the test set, and then summed to derive the final scores. “Medium” refers to the exchange reactions between the community and the environment, while “extracellular space” represents the exchange reactions between species and the community (D). There are significant differences in the flux of 12 metabolites, with an asterisk indicating a Wilcoxon test result with a p-value less than 0.001 (E).

Ultimately, the study identified several metabolites with significant impacts on hypertension, including choline(chol), 1-butanol(btoh), trimethylamine(tma), cytidine(cytd), betaine(glyb), methanethiol(ch4s), pyruvate(pyr), γ-aminobutyric acid(4abut), pyridoxamine(pydam), siroheme(sheme), L-serine(ser_L), and uracil(ura). Among these, three metabolites derived from dietary phosphatidylcholine: choline, trimethylamine N-oxide (TMAO), and betaine are already known to be associated with hypertension. Finally Wilcoxon tests were conducted on the 12 metabolites, with results showing p-values less than 0.001 for all (Figure 3E).

### 3.3 Alterations in Subnetworks within the Gut Microbial Interaction Network in Hypertension

The random walk algorithm was used to guild nodes in the differential GMiN, and identifying a total of 13 guilds(Figure 4A). Based on the shared 12 important metabolic flux-associated species identified by both the Random Forest and XGBoost models, species were ranked according to their frequency. Using the GSEA method, each guild was analyzed for enrichment of important metabolic flux-associated species. It was found that nodes in the third guild ranked significantly higher in terms of frequency among the important metabolic flux-associated species, with most of the rankings within the top 50(Figure 4B). In guild 3, there were severe disruptions in the interaction relationships at the genus level, particularly among species such as *Eubacterium, Ruminococcus, Klebsiella*, and *Parabacteroides*. These interactions showed significant differences. From the gutMDisorder database^40^, it was found that species within this guild, including *Bilophila, Desulfovibrio*, and *Ruminococcus*, have already been reported in two studies as being associated with hypertension. Moreover, in a recent study by Liping Zhao etal., it was found that the ratio of the ‘two competing guilds’ (TCG) of C1A and C1B is associated with various diseases, with 14 C1B members and 3 C1A members identified in KEPR (Figure 4C). Therefore, it is inferred that the altered interaction patterns within the KEPR may lead to abnormalities in the 12 important metabolites and ultimately influencing hypertension.

**Figure 4.**
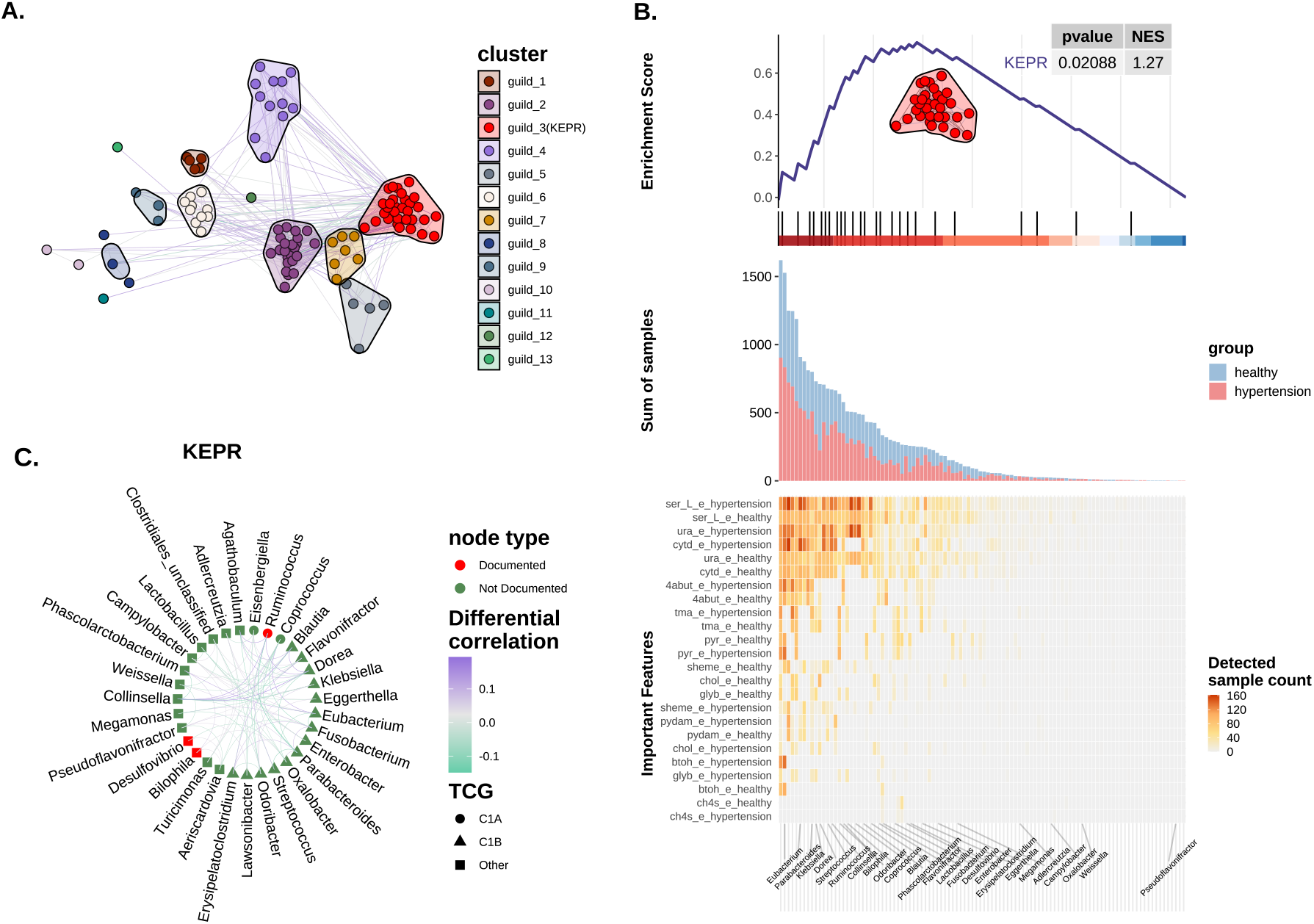
Alterations in Subnetworks within the Gut Microbial Interaction Network in Hypertension: The microbial correlation difference network was divided into 13 guilds using random walk algorithm (A). The GSEA algorithm was used to analyze the enrichment of the 12 important metabolites in microbial guilds. The 12 important metabolites are primarily associated with KEPR. NES is the normalized enrichment score. The color of the heatmap below represents the number of samples in which each species is associated with the 12 important metabolites (B).The microbial interactions in the KEPR have undergone significant changes and three species in the KEPR have already been shown to be associated with hypertension. The red nodes represent species that have already been proven to be associated with hypertension, green edges a strengthened negative correlationtions, and purple edges indicates a strengthened positive correlation (C).

### 3.4 Enhanced Mucin O-Glycan Degradation and Slower Acetic Acid Production Rate in the Intestines of Hypertensive Patients

#### 3.4.1 Enhanced O-glycan metabolism in the hypertensive GMFiN and important metabolite subnetworks

To uncover hypertension-related biomarkers at the metabolite level, metabolite correlation networks (GMFiN) were constructed (Figure 5A). To explore the differences in the topological structure of the overall GMFiN nodes. Based on four topological features: Betweenness, Closeness, Degree, and Eigen Centrality, the top 10 nodes with the most significant differences in topology were selected. It was found that the topological features of metabolites such as Nitrate(NO), Riboflavin(ribflv), Chorismate(chor), and Ursodeoxycholic acid(HC02194) were increased in the hypertension group. Additionally, the majority of released mucin-type O-glycan nodes exhibited lower eigen centrality in the hypertension samples (Figure 5B). But the degree of most mucin O-glycans and released mucin-type O-glycans increased in the hypertension network (Supplement Figure 1A). To investigate the differential sub-networks associated with the 12 important metabolites and the pathways involved, the differential sub-networks related to the 12 metabolites were extracted, and it was found that only 9 metabolites had significantly different edges with their neighboring nodes (Figure 5C). Pathway enrichment analysis was performed on the important metabolite sub-network nodes, and the results showed that O-glycan degradation and mucin-type O-glycan biosynthesis activities were enhanced in the hypertension network, while activities related to peptide metabolism, chloroalkane and chloroalkene degradation, and butyrate metabolism were reduced in hypertension. To further explore functional changes in the overall network, pathway enrichment analysis was performed on the top 100 nodes with the largest degree differences. The results showed that O-glycan degradation activity and mucin-type o-glycans biosynthesis was enhanced in the hypertension network, while peptide metabolism activity was reduced in hypertension (Figure 5C).

**Figure 5.**
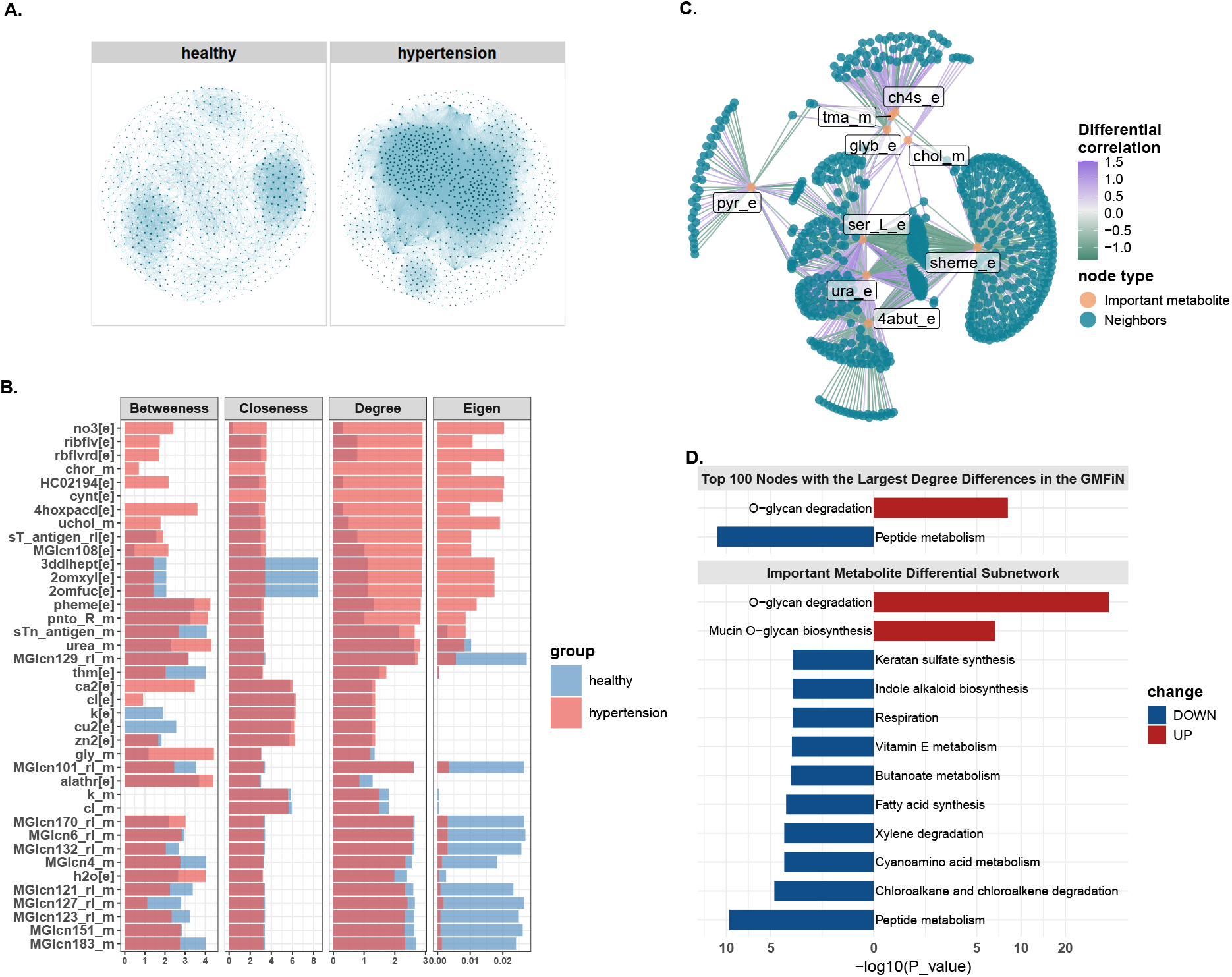
Enhanced Mucin O-Glycan Degradation and Slower Acetic Acid Production Rate in the Intestines of Hypertensive Patients: The metabolite networks of healthy individuals and hypertension patients were constructed (A). The topological structure of the metabolite correlation network nodes has undergone significant changes. The light red bars represent the topological values from the hypertension network, the light blue bars represent those from the healthy network, and the dark red areas indicate the overlapping values between the two (B). Among the 12 important metabolites, 9 metabolites exhibited significant differences in the edges of their correlation sub-networks. The color of the edges represents the magnitude of the differences, with green indicating a decrease in hypertension and purple indicating an increase. Orange nodes represent the important metabolites, and green nodes are neighboring nodes of the important metabolite, the edges connecting these nodes to the important metabolites show significant differences (C). Pathway enrichment was performed for the top 100 differential nodes in the metabolite correlation network as well as the nodes of the important metabolite sub-networks. Red indicates pathway upregulation in hypertension, while blue indicates pathway downregulation in the hypertension group (D).

### 3.4.2 Enhanced O-glycan degradation and reduced acetate and hydrogen sulfide production in the GMxN of hypertension

To explore the differences in microbial production and consumption of metabolites in the intestines of healthy individuals and hypertensive patients, cross-feeding networks were constructed. (Figure 6A). To explore the differences in the topological structure of the GMxN. Based on four topological features: Betweenness, Closeness, Degree, and Eigen Centrality, the top 10 nodes with the most significant differences in topology were selected. The topological structures of many common metabolites, such as water, hydrogen, and oxygen, were altered. Specifically, the topological feature values of metabolites like choline (chol), trimethylamine (tma), and betaine (glyb) were higher in healthy samples. In contrast, the in-degrees of metabolites such as hydrogen sulfide, siroheme (sheme), butyrate (but), Acetic acid (ac) were significantly lower in hypertension samples compared to healthy samples. This suggests that the more specieses in healthy samples produce choline, trimethylamine, betaine, hydrogen sulfide, sheme, butyrate and Acetic acid. Additionally, methanethiol (ch4s), nitric oxide (no) and Carob galactomannan(galmannan) were present only in the healthy samples (Figure 6B), and NO production was not detected in Campylobacter and Lactobacillus within guild 3(KEPR). This indicates that gut microbiota in healthy samples are capable of producing these three metabolites, whereas few gut microbiota in hypertensive patients produce them. Pathway enrichment analysis was performed on the top 100 nodes with the largest indegree differences in the GMxN. The results showed that microbial activity related to nucleotide interconversion, plant polysaccharide degradation, and O-glycan degradation was enhanced in the intestines of hypertensive patients, while activities related to pyruvate metabolism, methane metabolism, butanoate metabolism, and sulfur metabolism were reduced (Figure 6C). As products of mucin O-glycan degradation, some released mucin-type O-glycans are absorbed by gut microbes in healthy individuals, while in hypertensive patients, they are produced by gut microbes. At the same time, the absorption rate of some released mucin-type O-glycans decreases in hypertensive patients, whereas the absorption rate of mucin-type O-glycans increases. On the other hand, the rates at which gut microbes produce released mucin-type O-glycans and mucin-type O-glycans do not differ significantly, but the rate of Acetic acid (ac) production is downregulated in hypertensive patients (Figure 6D).

**Figure 6.**
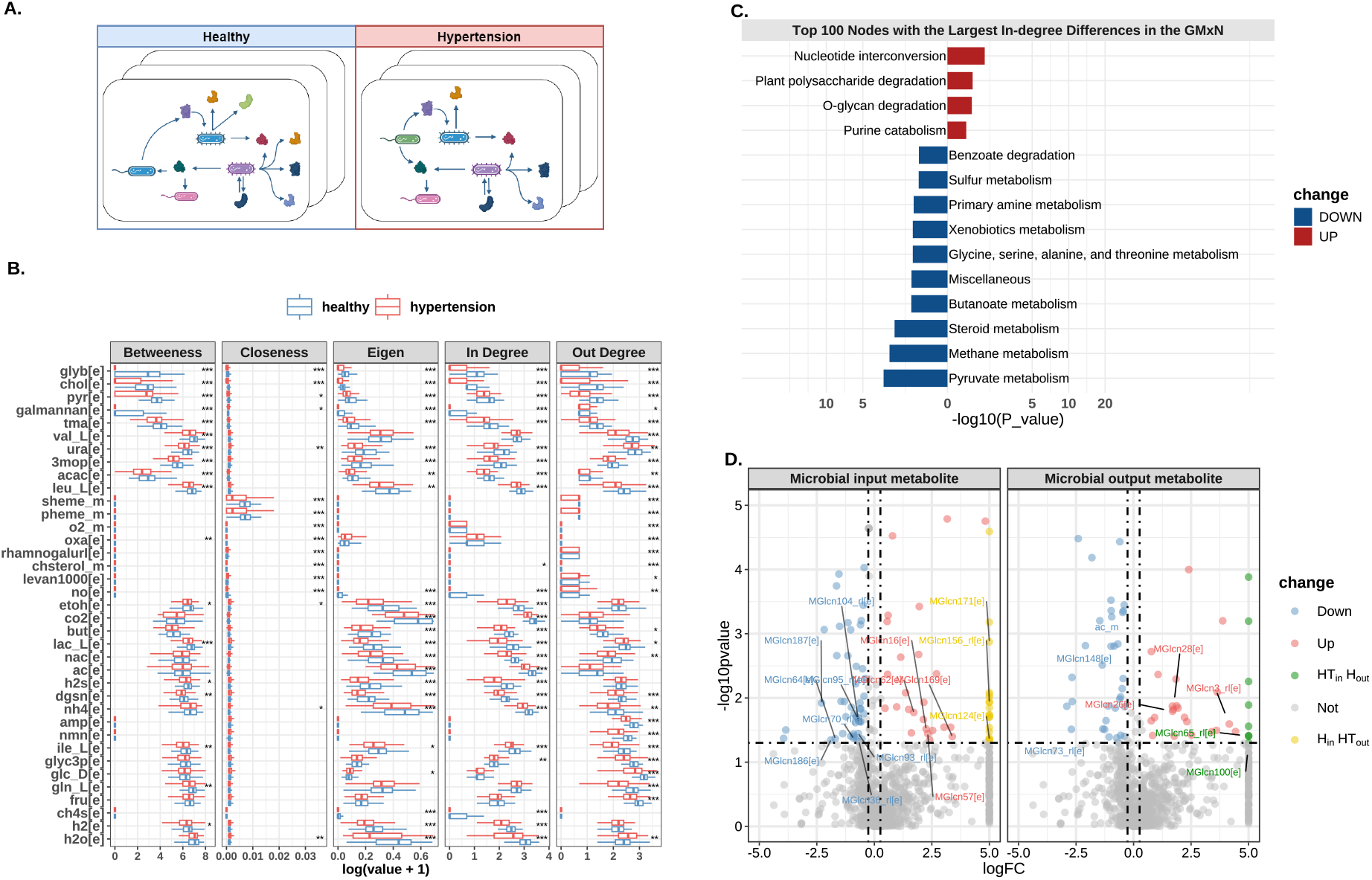
Enhanced O-glycan degradation and reduced acetate and hydrogen sulfide production in the GMxN of hypertension: Cross-feeding network diagram for each sample (A). Analysis of differences in the topological structure of the cross-feeding network, showcasing the top 10 metabolites with the most significant differences in each topological feature (B). Pathway enrichment was performed for the top 100 differential nodes in the Cross-feeding network (C). Analysis of the differences in overall consumption and production of metabolites, where red indicates upregulation, blue indicates downregulation, yellow signifies metabolites primarily output in healthy individuals but input in hypertensive individuals, and green indicates metabolites primarily input in healthy individuals but output in hypertensive individuals (D).

**Figure 7.**
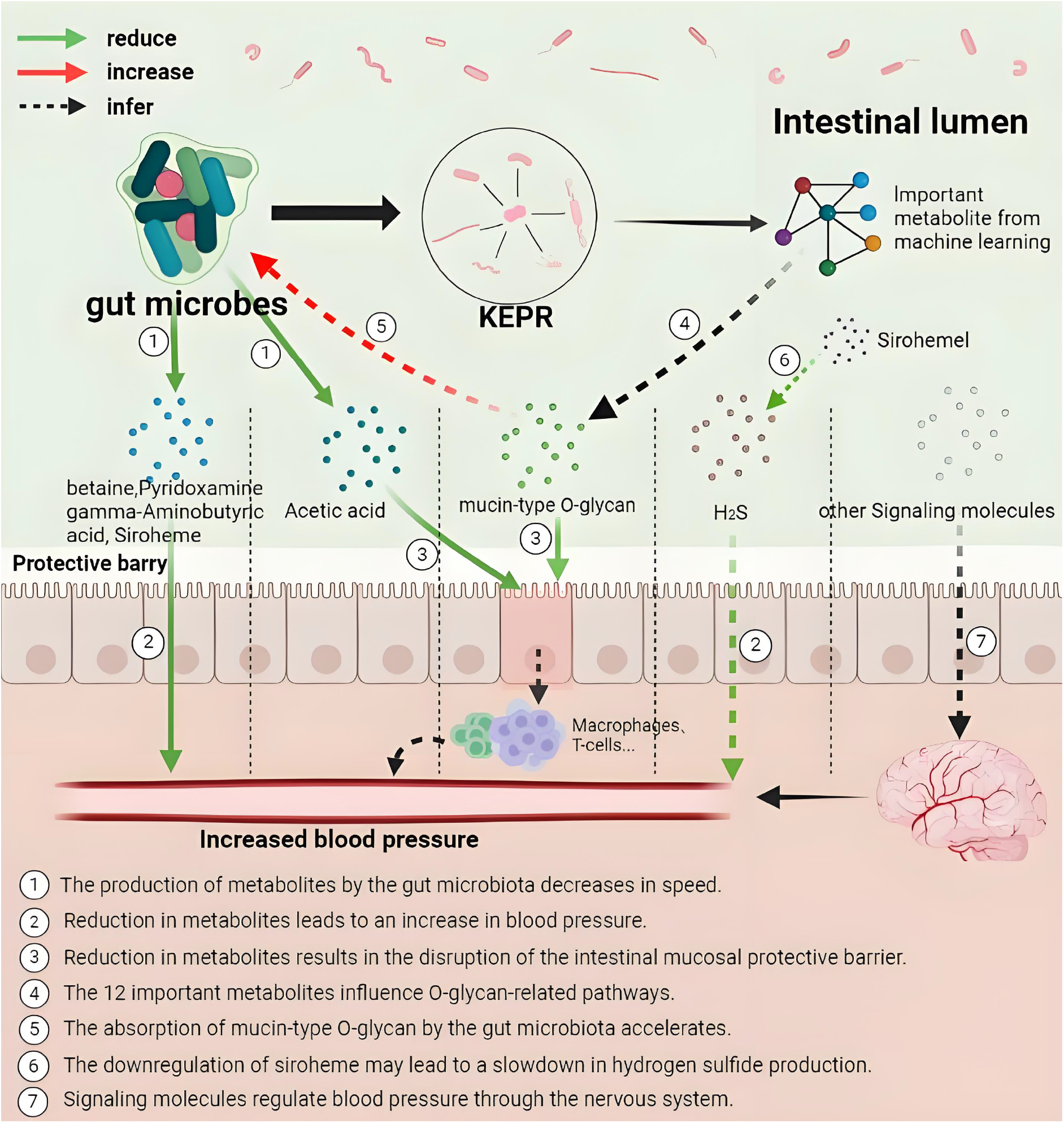
Hypothetical mechanisms of gut microbiota-mediated pathogenesis in hypertension inferred from gut microbiome data mining.

## 4 DISCUSSION

Hypertension remains one of the most significant risk factors for cardiovascular diseases (CVD), such as stroke and heart failure. It is the leading cause of non-communicable disease-related mortality, surpassing other major causes like smoking and metabolic disorders^55,56^. Currently, hypertension is one of the primary contributors to the global burden of disease, affecting over a billion adults, with its prevalence continuing to rise significantly^57,58^. Changes in the composition and function of the gut microbiota are closely linked to the onset and progression of hypertension^7,59^. Recent studies further suggest that the gut microbiota plays a critical role in the development of cardiovascular diseases, with its metabolic products being closely associated with these conditions^5^. However, the structure of the gut microbiota varies significantly between individuals due to diversity, leading to considerable differences in microbial abundance across different individuals, which poses challenges for microbiome research^60^. Gut microbiota data often exhibit high sparsity, meaning that only a small portion of species are detected in a sample, with many species having very low abundance or being entirely absent in certain samples.

To avoid the issues of significant inter-individual microbial differences and data sparsity, this study employs network-based approaches to indirectly explore hypertension-related microbes and metabolites through the study of metabolite interactions. Compared to traditional differential analysis methods like t-tests, Wilcoxon tests, and linear model-based approaches, constructing Random Forest and XGBoost tree-based models offers a better handling of complex feature interactions by identifying important features based on rules. Additionally, network construction focuses more on interactions and connections between nodes, allowing for the analysis of node importance through their topological characteristics.

Microbial interaction networks can reveal relationships between different microbial species, such as competition, cooperation, and mutualism. This aids in understanding the dynamic balance and ecological functions of microbial communities, shedding light on how these communities maintain stability under varying environmental conditions^61^. Network analysis can identify key microbes and metabolites that play crucial roles under specific environments or conditions. This is significant for developing microbial intervention strategies, optimizing fermentation processes, or improving human health. For example, in gut microbiota research, identifying key metabolic products can provide potential therapeutic targets^62,63^.

Metabolite network analysis aids in predicting the metabolic functions and pathways of microbial communities. By analyzing metabolic networks, it is possible to infer the pathways of production and consumption of specific metabolites, thus gaining insights into how metabolic activity of microorganisms influence the host’s physiological state. However, the construction of microbial and metabolite networks is based on statistical calculations, where edges are filtered solely based on significance p-values. The final cross-feeding network, being knowledge-based, further enhances the reliability of the experiment. Recent studies have demonstrated that gut microbiota play a crucial role in the development of cardiovascular diseases (CVD)^64^. The gut microbiota metabolizes dietary choline, phosphatidylcholine, and L-carnitine to produce trimethylamine (TMA), which is further oxidized to trimethylamine N-oxide (TMAO) in the liver^42,43^, a metabolite that promotes atherosclerosis. Betaine, derived from diet or choline oxidation, is associated with metabolic syndrome, dyslipidemia, and diabetes, and may be related to vascular diseases and other conditions^41^. In our analysis of metabolic flux data using Random Forest and XGBoost models, choline, trimethylamine, and betaine emerged as important features identified by both models, however there is not found abnormal in TMAO. Additionally, other common metabolites were also recognized, such as methanethiol, 1-butanol, pyridoxamine, gamma-aminobutyric acid (GABA), and siroheme. Existing research indicates that eight of these twelve metabolites may have potential roles in regulating blood pressure. In contrast, the other four are common metabolites associated with glycolysis and lack a direct relationship with hypertension (Table 1).

**Table 1.**
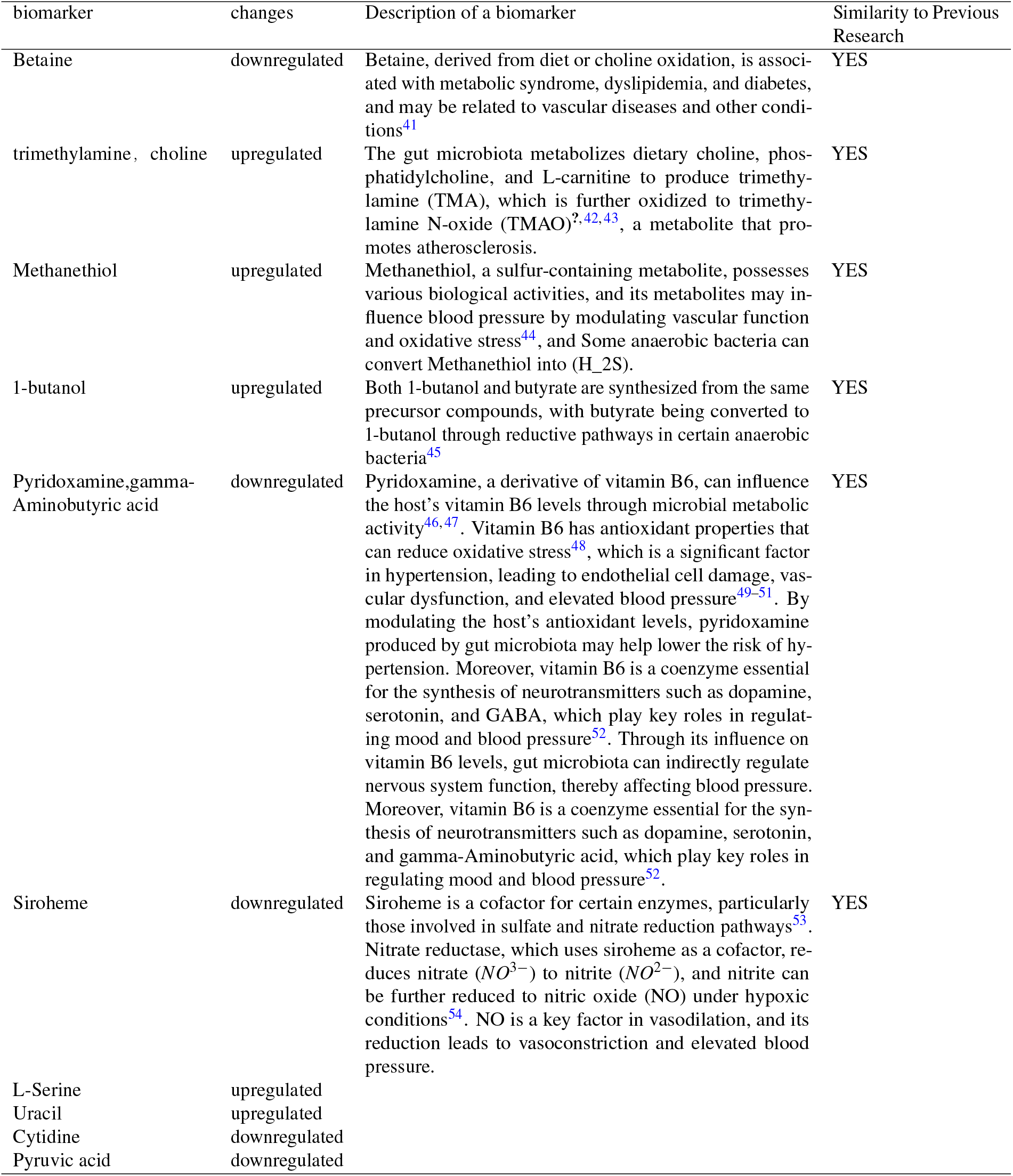
Hypertension-Related Metabolites Identified by Machine Learning.

The gut microbiota represents a complex system, and the construction of gut microbiota interaction networks (GMiN) or gut microbiota flux interaction networks (GMFiN) offers a more systematic approach to investigating the microorganisms or metabolites associated with the development of hypertension. In this study, differential GMiN were clustered using a random walk algorithm, and the enrichment of each guild with important metabolites was assessed using the Gene Set Enrichment Analysis (GSEA) algorithm. The results indicate that the subnetwork composed of species such as *Eubacterium, Ruminococcus, Klebsiella*, and *Parabacteroides* is disrupted and is associated with the dysregulation of these 12 metabolites which is linked to hypertension. Among these, three genus-level species: *Bilophila, Ruminococcus, and Desulfovibrio* were previously shown to be significantly downregulated in hypertensive samples in other studies^65,66^. Moreover, in a recent study by Liping Zhao etal., it was found that the ratio of the ‘two competing guilds’ (TCG) of C1A and C1B is associated with various diseases, with 14 C1B members and 3 C1A members identified in KEPR. the beneficial guild (C1As) comprises genomes rich in CAZy genes essential for the digestion of dietary fiber and genes instrumental in the production of SCFAs, such as butyrate, which play vital roles in nutrition, metabolism, immune function, and overall physiology. By contrast, what might be considered the detrimental counterpart (C1Bs) is characterized by genomes abundant with genes that confer antibiotic resistance and express VFs that may predispose to a pathogenic interaction with the host^67^. This indicates that a disturbance has occurred among most C1B species, which may further lead to hypertension in the host.

In the GMFiN, significant changes were observed in the network’s topological structure. Pathway enrichment analysis was performed on the top 100 nodes with the largest degree differences in the GMFiN. The results showed that O-glycan degradation and mucin-type O-glycan biosynthesis activity was upregulated in hypertensive patients, while peptide metabolism was downregulated. Among these 12 important metabolites, 9 metabolites exhibited significant differences in edge connections with neighboring nodes. It was found that O-glycan degradation activity and mucin-type O-glycan biosynthesis activity were elevated in the hypertension network, while activities related to peptide metabolism, chloroalkane and chloroalkene degradation, xylene degradation, fatty acid synthesis, and butanoate metabolism were reduced in the hypertension network.

In the GMxN, it was found that the topological characteristic values of choline (chol), trimethylamine (tma), and betaine (glyb) metabolites were higher in healthy samples, indicating that the metabolic activity of these metabolites in the GMxN is more active in healthy individuals. However, TMAO did not show noticeable changes. The metabolites methyl mercaptan (ch4s) and nitric oxide (NO) were found only in healthy samples, and NO production was not detected in KEPR’s *Campylobacter* and *Lactobacillus*, which produce more NO in healthy samples. This indicates that gut microbiota in healthy samples can transport these two metabolites, while hypertension patients’ gut microbiota almost cannot transport these metabolites. NO is a key factor in vasodilation, and its reduction leads to vasoconstriction and elevated blood pressure. Gut microbiota can influence host blood pressure regulation by producing NO and its precursors. NO activates soluble guanylate cyclase (sGC), increasing cyclic guanosine monophosphate (cGMP) levels, leading to smooth muscle relaxation, vascular dilation, and blood pressure reduction. Thus, NO plays an important role in the regulation of hypertension^68–70^. However, there was no significant difference in the final rate of NO production and consumption by the microbial community.

Methanethiol is produced by gut microbiota through the metabolism of sulfur-containing amino acids, such as methionine. In gut microbiota metabolism, Methanethiol can serve as a precursor for hydrogen sulfide *H*_2_*S* production. Some anaerobic bacteria can convert Methanethiol into *H*_2_*S*. The topological structure results from the GMxN show that the influx of *H*_2_*S* is significantly lower in hypertension compared to healthy individuals, indicating reduced production of *H*_2_*S* by gut microbiota in hypertension. *H*_2_*S* has been recognized as the third major gaseous signaling molecule, following nitric oxide (NO) and carbon monoxide (CO). This novel gaseous molecule has been shown to play a wide role in the regulation of various systems in the body^71^. Studies have found that *H*_2_*S* can directly induce relaxation of vascular smooth muscle by activating ATP-sensitive potassium channels(K_ATP channels) and inhibiting calcium channels, thereby promoting vasodilation^72^. An enrichment analysis of the top 100 differential in-degree nodes in the GMxN revealed that pathways such as O-glycan degradation and plant polysaccharide degradation were significantly upregulated in hypertension, while butyrate metabolism, pyruvate metabolism, methane metabolism, and sulfur metabolism were significantly downregulated.

O-glycans in the gastrointestinal tract serve as the first line of defense against external stimuli^73^. Glycosylation defects can impair mucin expression and disrupt the mucosal barrier, leading to microbial activation of caspase 1, interleukin IL-1β, IL-18, and other inflammasomes, which triggers inflammation and results in severe spontaneous bacterial-dependent colitis^74,75^. This inflammation can damage vascular endothelial cells, impairing their normal function. The inflammatory response generates a large amount of free radicals, leading to oxidative stress. Oxidative stress can further damage endothelial cells, causing endothelial dysfunction and contributing to increased blood pressure^76^. Therefore, the gut microbiota in hypertensive samples may affect the interaction patterns of mucin-type O-glycans with other metabolites, leading to metabolic dysregulation of mucin-type O-glycans and damage to the intestinal mucosa, which in turn affects the host’s blood pressure.

## 5 CONCLUSION

In this study, Random Forest and XGBoost models constructed using metabolic flux data jointly identified 12 metabolites associated with hypertension, including choline (chol), 1-butanol (btoh), trimethylamine (tma), cytidine (cytd), and betaine (glyb). These metabolic abnormalities were linked to disruptions in the KEPR guild species (*Eubacterium, Ruminococcus, Klebsiella, Parabacteroides* etc), and the dysregulation of these metabolites may lead to enhanced degradation and synthesis of mucin-type O-glycans, along with reduced butyrate metabolism activity. This could result in increased consumption of mucin O-glycans and decreased acetate production in the gut, potentially weakening the physical barrier that protects intestinal epithelial cells from pathogens and toxins. Damage to the intestinal mucosa can trigger inflammation, causing macrophages and T cells to release TNF-α, IL-6, and IL-1β, which activate endothelial cells, increase oxidative stress, and promote vasoconstriction. The large amount of free radicals generated in this process can further exacerbate oxidative stress, potentially leading to additional damage to vascular endothelial cells and causing endothelial dysfunction, which in turn leads to elevated blood pressure. Additionally, abnormalities in Siroheme may reduce or inhibit the secretion of metabolites or signaling molecules, such as microbe-derived hydrogen sulfide (*H*_2_*S*), which is critical for promoting vasodilation and lowering blood pressure. These changes may further contribute to the dysregulation of blood pressure control mechanisms.

## Supporting information

https://github.com/as147596/HT

## Author contributions statement

W.L.: Writing – original draft. Y.Z.: Writing – original draft. S.L.: Writing - original draft, data curation, validation, visualization. F.T.: Writing – review & editing, Methodology, Supervision, Validation, Investigation. L.D.: Funding acquisition, Writing – review & editing. F.Y.: Conceptualization, Funding acquisition, Project administration, Writing – review & editing, Writing – original draft. All authors reviewed the manuscript.

Must include all authors, identified by initials, for example: A.A. conceived the experiment(s), A.A. and B.A. conducted the experiment(s), C.A. and D.A. analysed the results. All authors reviewed the manuscript.

To include, in this order: **Accession codes** (where applicable); **Competing interests** (mandatory statement).

The corresponding author is responsible for submitting a competing interests statement on behalf of all authors of the paper.

This statement must be included in the submitted article file.

## Key Points

- The compositional nature of microbiome data is well-known but often neglected in the abundancebased statistical analysis.
- How best to classify highdimensional microbiome data remains an outstanding problem, partly due to the difficulty of correctly handling compositional data.
- The DBA-distal method selects the m

## SUPPLEMENTARY DATA

Supplementary data are available online at https://academic.oup.com/bib.

## ACKNOWLEDGEMENTS

Acknowledgements should be brief, and should not include thanks to anonymous referees and editors, or effusive comments. Grant or contribution numbers may be acknowledged.

## FUNDING

This research was financially supported by the National Natural Science Foundation of China [62102065], Joint Funds for the Innovation of Science and Technology, Fujian province (Grant number: 2022J05055), Fujian Medical University Research Foundation of Talented Scholars [XRCZX2022003].

## CONFLICT OF INTEREST

The authors declare that the research was conducted in the absence of any commercial or financial relationships that could be construed as a potential conflict of interest.

## DATA AVAILABILITY

The data and code used in this study are available on GitHub at https://github.com/as147596/HT

## Notes

https://github.com/as147596/HT

### Competing Interest Statement

The authors have declared no competing interest.

### Summary of Updates

Beautified the images and formatting, and added some discussions.

