## Supplementary figures and images for "Gut microbial metabolic flux disorder in hypertension"

### https://github.com/as147596/HT

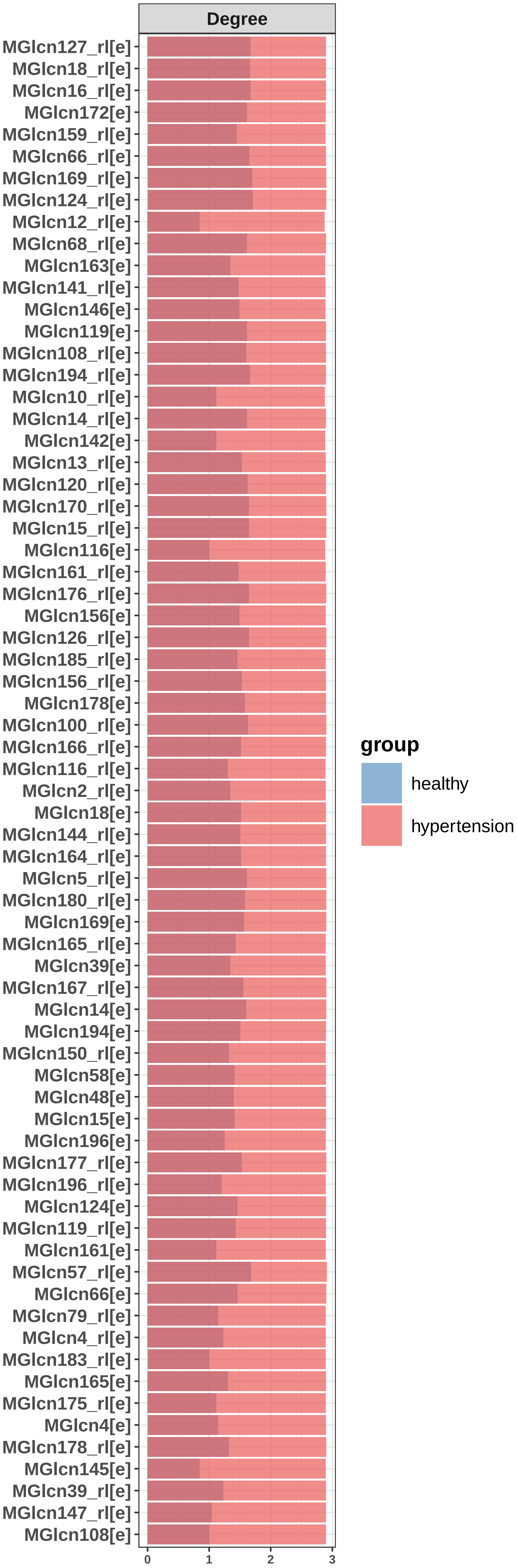
